# Exploring the Therapeutic Potential of Cannabis Constituents in Parkinson’s Disease: Insights from Molecular Docking Studies

**DOI:** 10.1101/2023.11.11.566677

**Authors:** Moawaz Aziz, Hafsa Rehman, Azhar Iqbal, Allah Nawaz, Momina Hussain, Tehmina Siddique, Sheikh Arslan Ashraf Sehgal, Muhammad Sajid

## Abstract

Cannabis, often known as marihuana, marijuana, hashish, and hash, belongs to the genus Cannabis sativa L. This plant has excellent potential for the treatment of several brain disorders. Phytochemical compounds in this plant act as antioxidants, preserving synaptic plasticity and preventing neuronal degeneration. The neurodegenerative condition Parkinson’s has emerged as one of the most significant health concerns of the twenty-first century. A detailed in silico molecular docking study was carried out to assess the neuroprotective effects of cannabis compounds against four potential targets of PD, including monoamine oxidase B (MAO-B), catechol-O-methyltransferase (COMT), alpha-synuclein (ASN), and Adenosine A2A receptor (A2A). Physicochemical properties, drug-likeness, toxicity, and ADMET profiles were also investigated. In this docking study, the cannabis compound cannabicyclol showed a superior docking score of −10.8 kcal/mol with the MAO-B protein. Based on these results, cannabicyclol and the target protein MAO-B were used to perform MD simulations to analyze their stability at 100 ns. Furthermore, it is crucial to carry out in vitro and in vivo investigations to enhance the potency of cannabis components and understand the processes underlying the suppression of Parkinson’s disease-related enzymes.

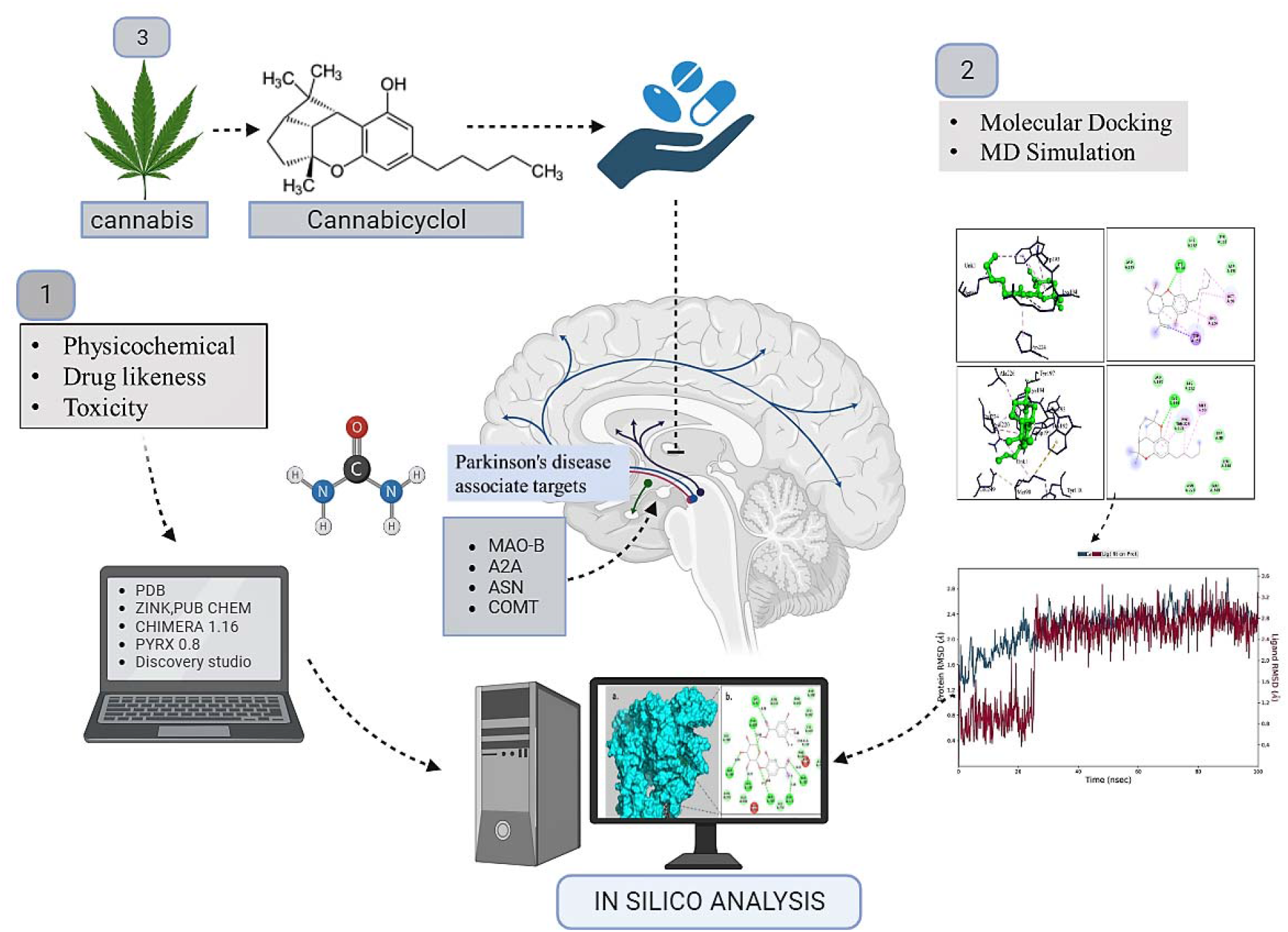

## Introduction

Cannabis sativa L. (C. sativa) is a flowering, dioecious, cross-pollinated plant with a height range of one–two meters. Common names for Cannabis sativa include hemp, cannabis, and marijuana [1]. It has been used as a medicinal plant in many regions of the Middle East and Asia since the 6th century BC [2]. Cannabis sativa is known for its medicinal and psychoactive properties. It can be used for recreational, therapeutic, and industrial purposes [3]. Currently, many studies are being conducted on cannabis because of its unique phytochemical components or secondary metabolites. The most active cannabinoid metabolites, phytocannabinoids, are found in the glandular trichomes of the female plant bark and mainly in the leaves [4, 5]. The cannabis plant contains approximately 565 known cannabis compounds, and more than 150 substances are categorized as phytocannabinoids [6–8]. Owing to their potent antioxidant, anti-inflammatory, and neuroactive effects, they have attracted much interest from researchers. It can cure several neurodegenerative illnesses [9].

CBD and THC are two types of cannabinoids that interact with the endocannabinoid system (ECS). These cannabinoids have properties that may help in the treatment of various diseases, including Alzheimer’s disease, amyloid-β- and NFT-related diseases, and neurogenesis [10]. The most prevalent substance in cannabis is Tetrahydrocannabinol (THC). The psychoactive potential of THC in cannabis has increased people’s curiosity. The substances in cannabis preparations used for psychoactive purposes are receptors in the endocannabinoid system (ECS) [11]. Additionally, the active ingredients of cannabis help regulate inflammatory disorders and cancer-related symptoms [12]. Owing to their sedative and relaxing properties, the most recent therapeutic application of these substances’ entails researching their potential for controlling sleep problems, particularly insomnia [13]. Cannabis has been indicated as an alternative treatment for many diseases including inflammatory bowel disorders (IBDs), glaucoma, sclerosis multiplex (SM), multiple sclerosis, schizophrenia, anxiety, Alzheimer’s disease, chronic pain, Parkinson’s disease, Huntington’s disease, and COVID-19 Table 1 [14–17].

**Table 1.**
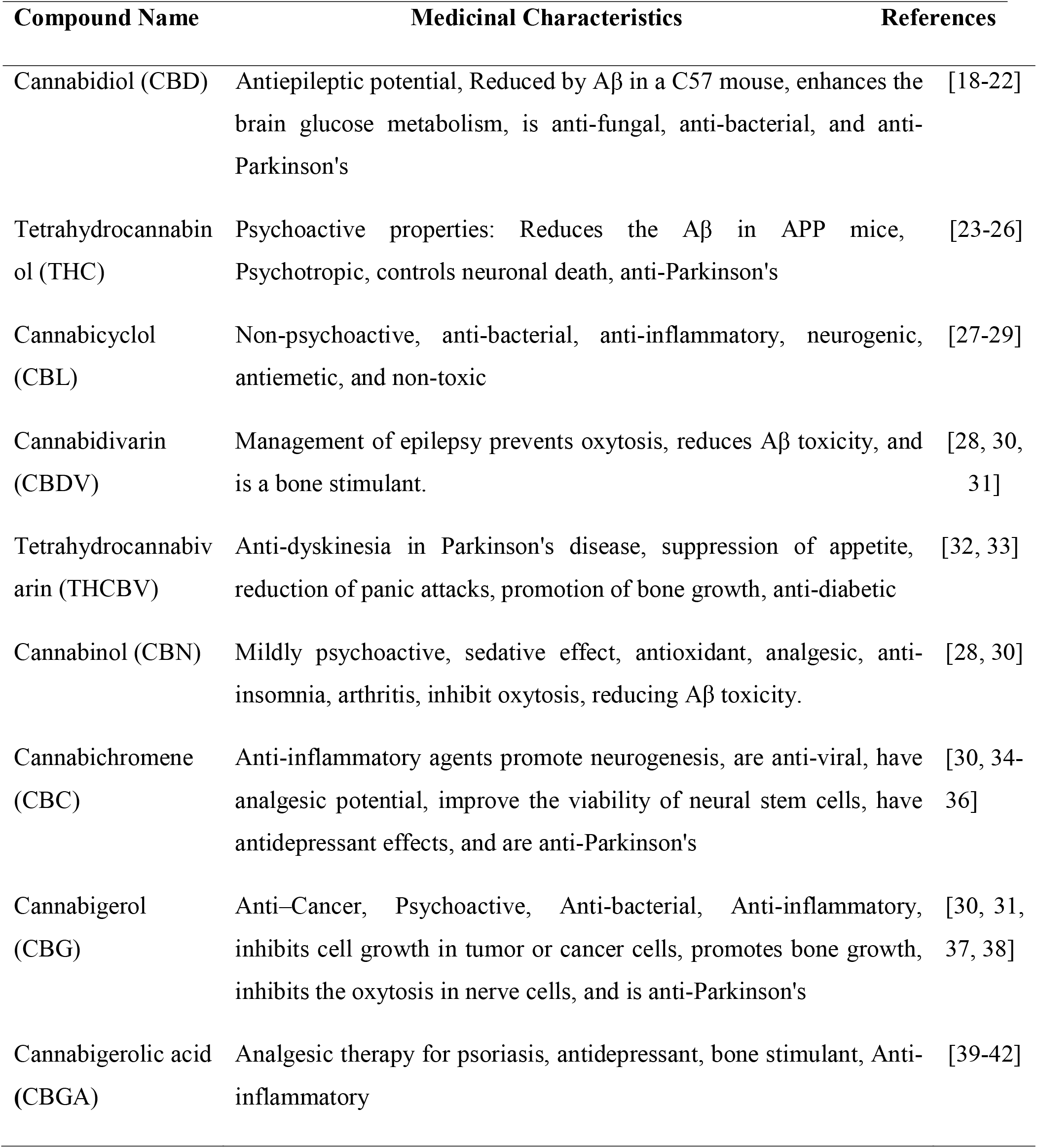
Primary metabolites reported in C. sativa and their therapeutic properties.

Other naturally occurring cannabinoids produced from Cannabis sativa are categorized into a variety of groups, including **Cannabidiol** (CBD), **Cannabichromene** (CBC), **Cannabicyclol** (CBL), **Cannabitriol** (CBT), **Cannabielsoin** (CBE), **Cannabinol** (CBN), and other unconventional types [43]. Cannabinoids interact with several receptors to exert their wide range of beneficial biological effects in the central nervous system and modulate the toxicity of AD-associated proteins. They comprise many non-cannabinoid receptors, such as G-protein protein complexes (GPR55 and GPR3) and ion channels, as well as the cannabinoid-binding sites CB1 and CB2 [44].

Parkinson’s disease (PD) is the most prevalent neurodegenerative disease that affects the nigrostriatal pathway in the brain as a result of byproduct development (Lewy bodies) and loss of dopamine [45]. This can lead to difficulty in moving forward and making decisions. This pathway is responsible for the transport of dopamine, a neurotransmitter that is then used to control movement. Due to a lack of dopamine, people with Parkinson’s disease often have difficulty moving their limbs and muscles [46]. Some people with PD has decreased numbers of dopaminergic cells in the substantia nigra, a brain region. Most of those with this deficit were aged over 60 [47]. Many medications are used to treat Parkinson’s disease (PD). Carbidopa, levodopa, dopamine agonists, MOA-B inhibitors, COM inhibitors, and amantadine are all used to treat PD [48, 49]. However, these drugs have many adverse effects. Some of these complications may lead to further complications [50]. We need to identify new drugs with the fewest side effects and determine the most effective drugs. Many active phytochemicals can significantly counter both Alzheimer’s and Parkinson’s diseases.

Alpha-synuclein significantly influences the progression of Parkinson’s disease. This protein is prevalent in the presynaptic terminals and is widely distributed throughout the brain [51]. Intractable clumps, known as Lewy bodies, are characteristic of Parkinson’s disease and are formed when alpha-synuclein aggregates [52]. The accumulation of α-synuclein (α-syn) in inclusions (Lewy bodies) and dysregulation of dopamine signaling pathways play a direct role in the pathophysiology of Parkinson’s. A potential treatment for Parkinson’s disease (PD) involves preventing alpha-synuclein aggregation [53]. Alpha-synuclein inhibitors have shown promise in preclinical research for lowering alpha-synuclein aggregation, enhancing motor symptoms, and safeguarding dopaminergic neurons [54]. These inhibitors may also prevent other alpha-synuclein-related degenerative processes such as mitochondrial failure and neuroinflammation [55]. Additional studies are required to ascertain the effectiveness, safety, and ideal dosage of alpha-synuclein inhibitors for treating PD. The creation of potent and specific inhibitors can control PD and possibly stop PD progression [56].

Monoamine oxidase (MAO) is an enzyme that breaks down the neurotransmitters. There are two different isoforms of MAO-A and MAO-B. MAO-A is primarily found in the brain, whereas MAO-B is more widely distributed throughout the body. Dopamine is oxidatively deaminated in the striatum by mitochondrial flavoenzyme monoamine oxidase B (MAO-B). MAO-B is a primary target of pharmacological interventions for Parkinson’s [57]. As you age, your brain has more MAO-B, which means that you produce more dopamine. This may increase dopamine metabolism and hydrogen peroxide production, which can result in neuronal cell death [57]. Therefore, inhibition of MAO-B is a promising therapeutic strategy for treating Parkinson’s disease.

Adenosine receptor (AR) antagonists have recently attracted much interest in the research and development of non-dopaminergic medicines for treating Parkinson’s disease (PD) [58]. Among the four human adenosine receptors (A1, A2A, A2B, and A3), the A2A receptor has received the most attention because it can be used to treat Parkinson’s disease [59]. A2AAR antagonists have been investigated as a possible class of non-dopaminergic antiparkinsonian drugs that might help patients with signs of depression (a non-motor PD symptom) [60]. In the forced swim and tail suspension tests, the A2AAR antagonist KW-6002 was reported to have antidepressant action in mice [61, 62]. Epidemiological and practical studies have shown that adenosine A2A receptor antagonists have neuroprotective benefits [63].

Human COMT is expressed in two different isoforms: membrane-bound COMT (MBCOMT), the most prevalent isoform in the brain, and the soluble form (SCOMT). Catechol-O-methyltransferase catalyzes the transfer of a methyl group from the cofactor S-adenosylmethionine (SAM) to oxygen hydroxylate the substrate catechol or substituted catechol [64]. COMT breaks down dopamine in the brain, especially in the prefrontal cortex and basal ganglia [65]. An essential component of the process through which COMT affects Parkinson’s is the regulation of dopamine levels [66]. Parkinson’s disease causes a significant loss of dopamine-producing neurons in the substantia nigra, which lowers dopamine levels in the brain [67]. By preventing dopamine from being broken down, COMT inhibition can increase dopamine levels [68]. This improves dopaminergic transmission and may help with the neurological symptoms of Parkinson’s [69].

The objective of this study was to conduct a comprehensive in silico molecular docking study and molecular dynamic simulation to analyze how different cannabis constituents might help in the treatment of Parkinson’s. Furthermore, physicochemical properties, drug-likeness, toxicity, and ADMET profiles were examined to confirm their safety and effectiveness in treating PD.

## MATERIALS AND METHODS

### Selection and Ligand preparation

Fifteen cannabis compounds with medicinal properties were selected from available literature and databases. The compounds were selected based on their reported medicinal potential in the available research. The *PubChem* and *Zinc databases* were used to find the chemical structures of the selected substances: https://zinc.docking.org/substances/home/. The selective 15 cannabis compounds were downloaded from PubChem and the ZINC database in SDF format. After retrieval, the energy of the compounds was minimized using *Chimera 1.16*.

### Retrieval and Protein preparation

The structure of Parkinson’s disease-associated proteins A2A receptor with PDB I’d 3EML, COMT with PDB I’d 1H1D, MAO B with PDB I’d (2C65), and ASN with PDB I’d (1XQ8) have been retrieved from the *Protein Data Bank* https://www.rcsb.org/. The ligands and water molecules were removed from the crystal structures of the selected enzymes. The missing hydrogen atoms were then added by editing and minimizing the structure using *Chimera 1.16*.

### ADME/T prediction

The ADMET characteristics were predicted earlier to reduce the compound’s failure rate for future processes. Basic Pharmacokinetic and ADMET characteristics of natural chemicals such as molecular weight, log P, hydrogen bond donors, hydrogen bond acceptors, topological polar surface area, number of rotatable bonds, and Lipinski’s Violations were calculated by *swissADME* and *pkCSM* https://biosig.lab.uq.edu.au/pkcsm/ [74]. The essential information that these projections offer is the oral absorption, tissue distribution, metabolic stability, and possible toxicological concerns of the substance. This procedure assists in the discovery of substances with favorable pharmacokinetic characteristics.

### Toxicity forecasting

The ProTox-II database was used for toxicity forecasting to evaluate the possible toxicities of the selected chemicals. We predicted the toxicity of the substances by entering the smiles of compounds into *ProTox-II*, concentrating on the LD50 (median fatal dosage) values, and assigning toxicity classes https://tox-new.charite.de/protox_II/. This process facilitates the selection of safer treatment options by identifying chemicals with lower predicted toxicity.

### Biological Activity Forecasts of Cannabicyclol

The *PASS* website https://www.way2drug.com/PassOnline/ was used to forecast the Biological Activity of the Cannabicyclol by using SMILES from PubChem. By examining its biological activity spectrum, we learned more about the substance’s possible mechanisms of action and its applicability as a treatment for Alzheimer’s disease. This was done using MNA descriptors, suggesting that a compound’s chemical makeup influences its biological behavior [73].

### Target Ligand Docking

The Lamarckian genetic approach was used to perform a molecular docking study using the *PyRx-v0.8* virtual screening tool https://pyrx.sourceforge.io/. All ligands were docked separately to the target proteins. First, the enzyme was loaded as a macromolecule and 15 cannabis ligands were loaded individually. The grid box was in the middle of the enzyme’s binding location to provide the best conformational docking results, and the grid was placed at the Parkinson’s disease-associated targeted enzymes. The grid position was set at (x: 238.437, y: 80.77, z: 13.60) for Alpha-synuclein, at (x: 53.9113, y: 149.558, z: 17.1728) for MAO-B, at (x: 0.889), y: 2.9022, z: 49.5469) for Adenosine A2A Receptors, and at (x: 1.3135, y: 8.5065, z: −44.6610) for COMT. BIOVIA Discovery Studio v. 21.1.0 was used to visualize the 2D and 3D structures of the interacting residues of enzymes with ligands. This helped to determine the types of amino acids involved in binding, how they interact, and which types of bonds are formed between proteins and selected ligands. Deprenyl, Rasagiline, and selegiline are the MAO-B inhibitors that displayed binding affinity −7.3 kcal/mol, −7.9 kcal/mol, and −7.3 kcal/mol, respectively, were chosen as baselines to assess the efficacy of MAO-B receptor analogs [74]. Methylphenidate and one methyl-4-phenyl-1,2,3,6-tetrahydropyridine, having binding affinities of −5.1 kcal/mol and 4.7 kcal/mol, were chosen as baselines to assess the efficacy of the alpha-synuclein receptor analogs. Levodopa is the A2A receptor inhibitor with binding affinities of −6.6 kcal/mol chosen as a baseline to assess the efficacy of Adenosine A2A receptor analogs, and entacapone and tolcapone, having binding affinities of −5.4 kcal/mol and −6.9 kcal/mol, was chosen as baselines to assess the efficacy of COMT analogs.

### MD Simulation

The *Desmond 2023-3* version was used for the molecular dynamic simulation https://www.schrodinger.com/products/desmond. In this study, we selected the MAO-B protein and cannabicyclol molecule with the highest molecular docking score for molecular dynamics (MD) simulation analysis. We ran the simulation at 310 Kelvin for 100 ns. An orthorhombic box with dimensions of 10 mm × 10 mm × 10 mm was generated. The forcefield Optimized Potentials for Liquid Simulations (OPLS)-3e was applied to the single point charge (SPC) water model to solve the system. In total, 53,853 atoms and 15,255 water molecules were present in the system. To maintain stability, we applied an ensemble known as NPT, combined with a Nose-Hoover thermostat and barostat. We generated graphs presenting the root mean square deviation (RMSD) and root mean square fluctuation (RMSF), which helped us understand how the shape of the complex changes with time and better understand its behavior.

**Table 2.**
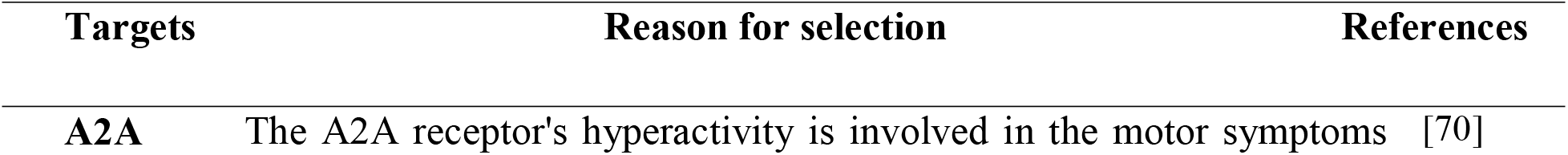
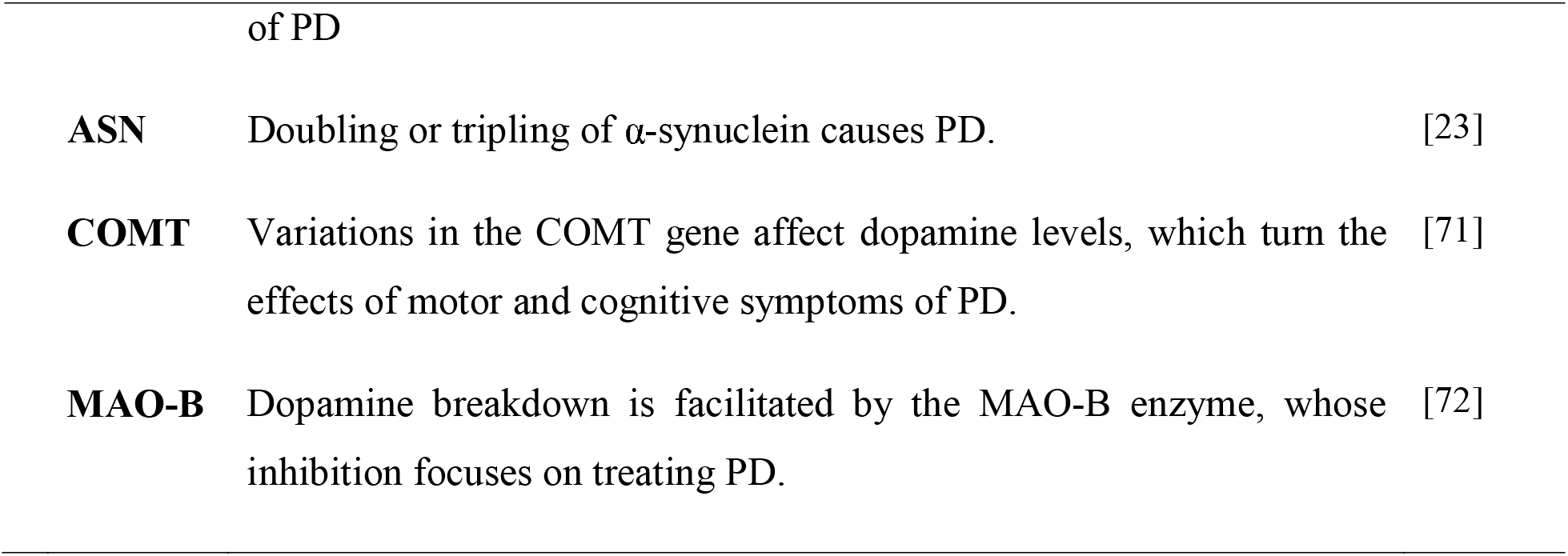
Targets in PD.

| Targets | Reason for selection | References |
| --- | --- | --- |
| A2A | The A2A receptor's hyperactivity is involved in the motor symptoms | [70] |
| **ASN** | Doubling or tripling of α-synuclein causes PD. | [23] |
| **COMT** | Variations in the COMT gene affect dopamine levels, which turn the effects of motor and cognitive symptoms of PD. | [71] |
| **MAO-B** | Dopamine breakdown is facilitated by the MAO-B enzyme, whose inhibition focuses on treating PD. | [72] |

## Results and Discussion

### Basic physicochemical premises of cannabis Compounds

Drug development is a lengthy and expensive process. Computational methods can predict the elements that improve the performance of composites. Lipinski’s rule of five was used to determine the pharmacokinetic characteristics of the cannabis constituents, which are potential natural lead molecules for new drugs. A few drug-relevant characteristics, including Molecular Weight, logP, Rotatable Bonds, H-Bond Acceptor, H-bond donor, and Topological Molecular Polar Surface Area, are shown in Table 3. Only one violation existed for certain compounds. All the data mentioned earlier show that these substances have great promise as drug-like molecules and therapeutic agents for various disorders, including neurodegenerative diseases.

**Table 3.**
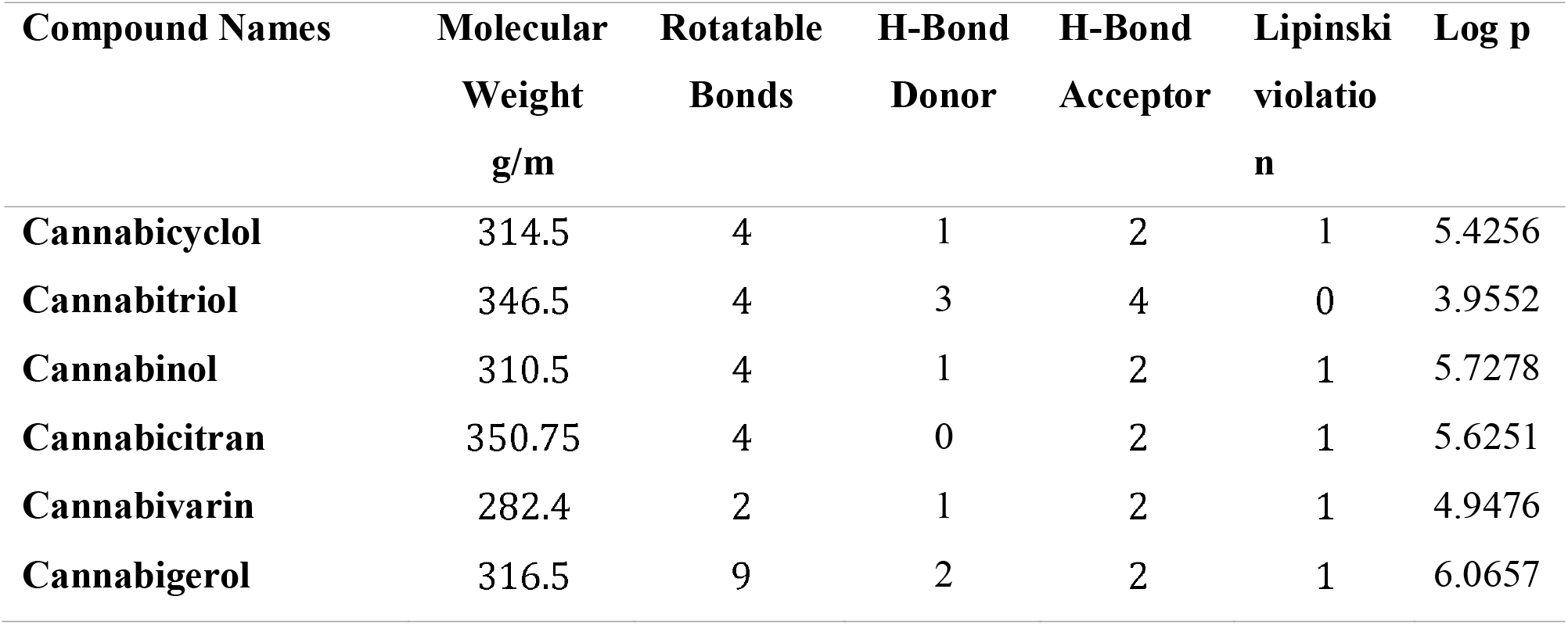
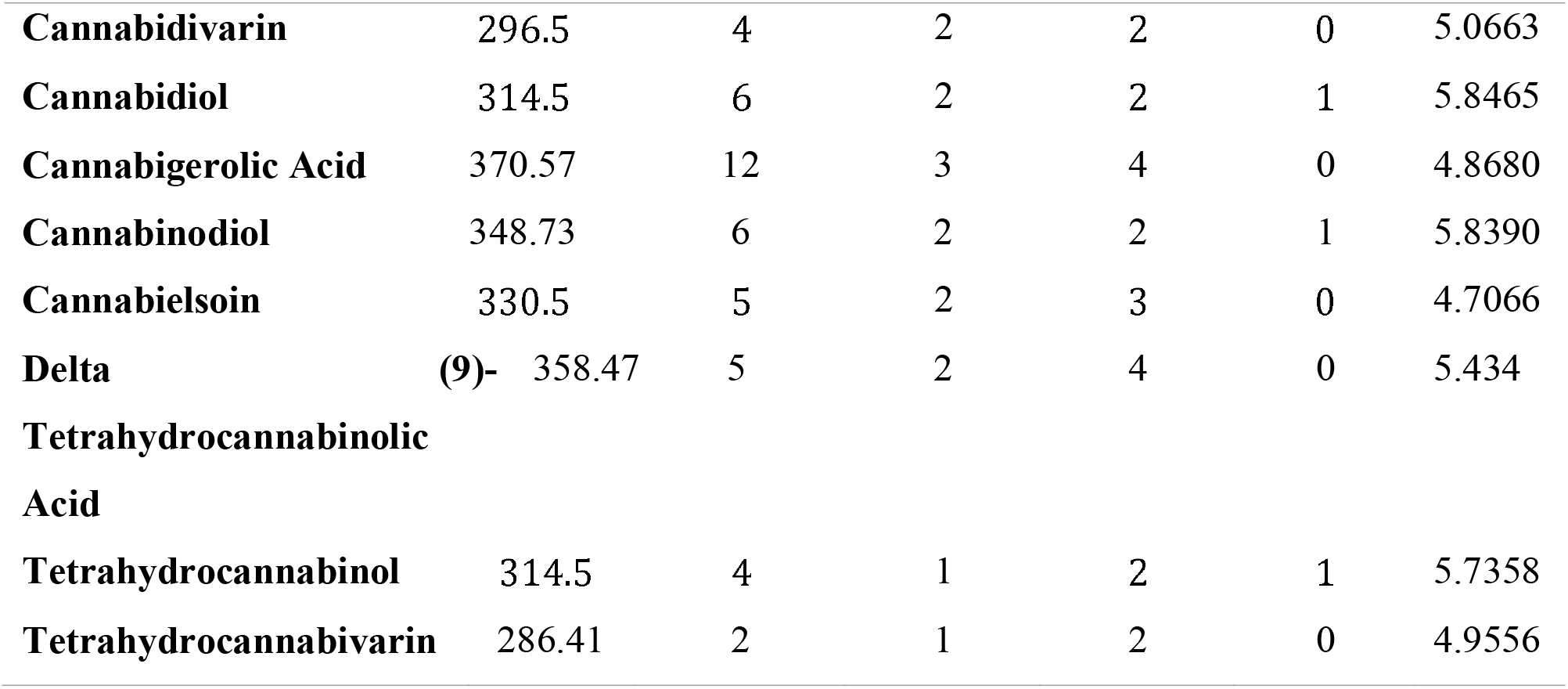
The physicochemical properties of 15 selective compounds.

### ADMET and Toxicity Appraisal

PkCSM and the SWISS-ADME online web server predicted the ADMET values for all constituents. It has characteristics such as BBB permeability, GI absorption, CNS permeability, and CYP2D6 inhibitory activity (Table 4). The predicted LD50 for acute oral toxicity varied from cannabinol (13500 mg/kg) to tetrahydrocannabinol (482 mg/kg). These substances have LD50 values in class IV and above, indicating that they are non-toxic if ingested.

**Table 4.**
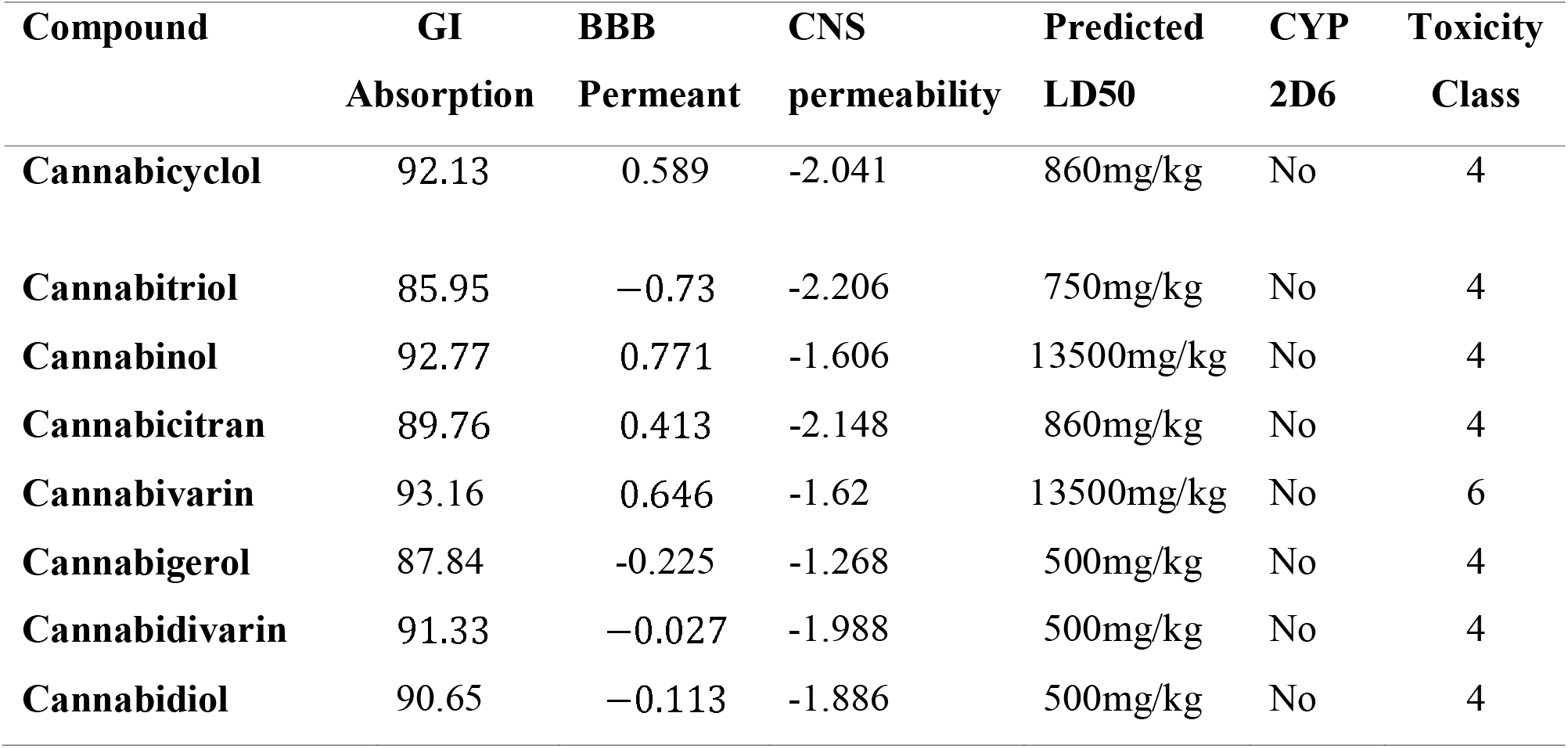
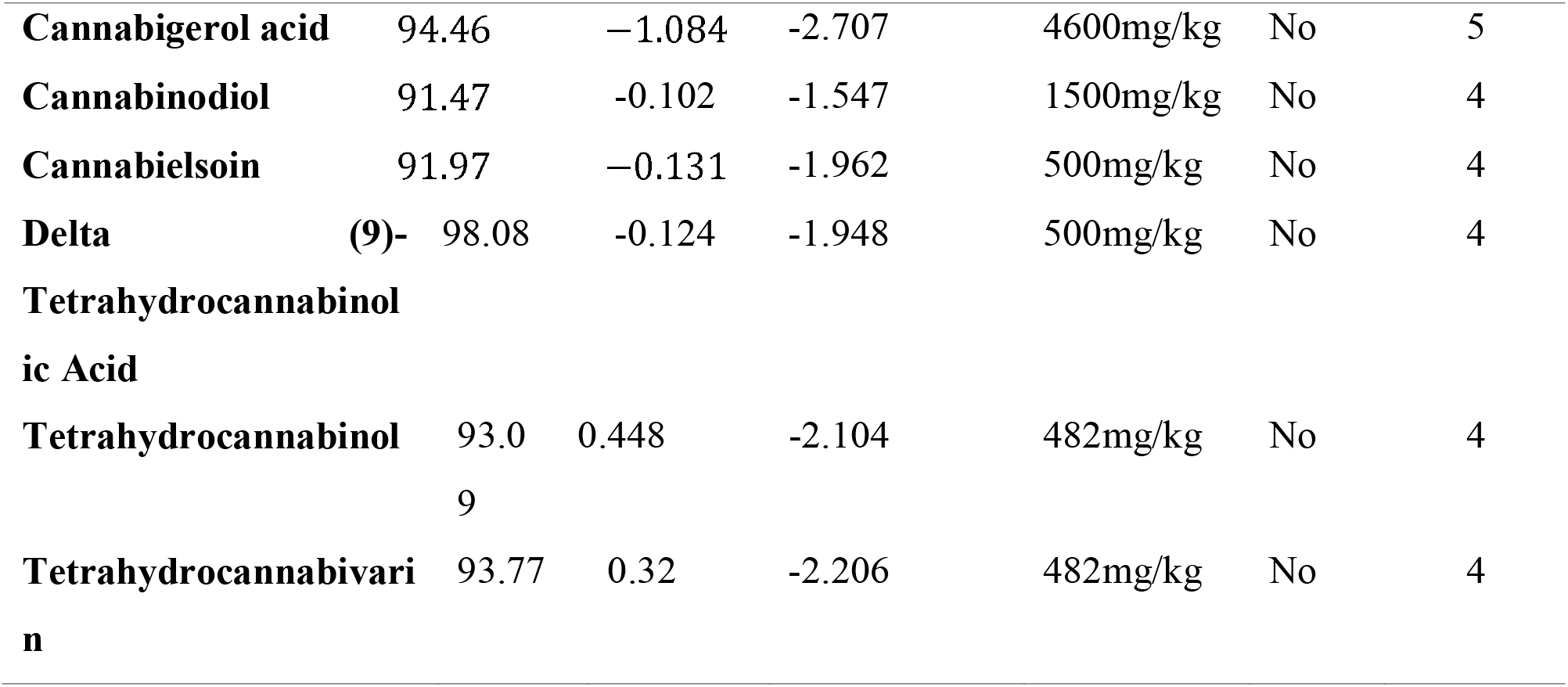
ADMET parameters and toxicity prediction of the compounds.

### Bioactive Spectra Prediction for Cannabicyclol

To determine the ten desired biological activities, the prediction of activity spectra for substances (PASS prediction) was carried out for cannabicyclol. In PASS predictions, the probabilities Pa (for “to be active”) and Pi (for “to be inactive”) are complementary. The technique attempts to select compounds with a higher chance of activity and a lower likelihood of inactivity by assessing the balance between Pa and Pi. This strategy aids in the discovery of exciting compounds with accurate predictions of biological activity. These biological activities suggest that these compounds are advantageous for treating neurodegenerative diseases. The test results for the selected ligands are listed in Table 5.

**Table 5.** The PASS prediction results display the biological functions of the top-performing docked ligand.

| **Biological activities** | **Pa** | **Pi** |
| --- | --- | --- |
| Neurotrophic factor enhancer | 0,580 | 0,003 |
| Immunosuppressant | 0,571 | 0,033 |
| Neuromuscular inhibiting agent for acetylcholine | 0,496 | 0,077 |
| Cytoprotectant | 0,421 | 0,088 |
| Chemopreventive | 0,410 | 0,021 |
| Treatment for acute neurological conditions | 0,405 | 0,139 |
| Kinase inhibitor | 0,394 | 0,069 |
| Anti-inflammatory | 0,242 | 0,222 |
| Protein synthesis blocker | 0,184 | 0,062 |
| TP53 expression enhancer | 0,622 | 0,043 |

### Molecular Docking appraisal

The docking score was always displayed negatively in the AutoDock program, with a more significant negative number indicating more potency. All 15 cannabis ligand molecules were successfully docked to Alpha-synuclein, MAO-B, A2A, and COMT. Given the comparison between lower binding energy and higher affinity, molecules with the most significant reduction in binding energy were deemed the best compounds for suppressing the protein target. The four best ligands for each target were selected from among the 15 ligand molecules, based on their affinity for binding to the designated targets. The binding affinities of cannabis constituents to the three selected target enzymes of Alzheimer’s disease are presented in Table 6. The docking scores ranged from 4.7 for alpha-synuclein 10.8 and MAO-B kcal/mol.

**Table 6.** Binding affinities (kcal/mol) of Cannabis compounds against selected target proteins.

| Compounds | A2A | COMT | MAO-B | Alpha-synuclein |
| --- | --- | --- | --- | --- |
| **Cannabicyclol** | −10.6 | −6.4 | −10.8 | −6.4 |
| **Cannabitriol** | −8.5 | −5.6 | −7.7 | −6.2 |
| **Cannabinol** | − 9.7 | −6.4 | −9.6 | −5.6 |
| **Cannabicitran** | −8.5 | −6.1 | −9.1 | −6.1 |
| **Cannabivarin** | −8.4 | −6.0 | −9.6 | −5.7 |
| **Cannabigerol** | −7.6 | −5.7 | −9.1 | −4.7 |
| **Cannabidivarin** | -6.4 | -5.7 | -9.1 | -5.7 |
| **Cannabidiol** | -7.5 | -5.2 | -8.4 | -5.5 |
| **Cannabigerolic acid** | -8.4 | -5.5 | -9.4 | -4.8 |
| **Cannabidiol** | -7.1 | -5.5 | -8.5 | -5.3 |
| **Cannabielsoin** | -7.5 | - 6.1 | -5.8 | -5.7 |
| **Delta (9)-**<br>**Tetrahydrocannabinolic**<br>**Acid** | -8.0 | -5.8 | -8.9 | -5.9 |
| **Tetrahydrocannabinol** | -8.5 | -6.0 | -9.3 | -5.8 |
| **Tetrahydrocannabivarin** | -8.5 | -5.8 | -9.2 | -6.0 |

### Molecular interaction of the MAO-B active residues with the highest docking score constituents

Ligands cannabicyclol, Cannabigerolic acid, tetrahydrocannabivarin, and Cannabidivarin had the highest scores of −10.8 kcal/mol, −9.4 kcal/mol, −9.2 kcal/mol, and −9.6 kcal/mol with the MAO-B protein, as shown below. Three substances, Cannabicyclol, Cannabigerolic acid, and tetrahydrocannabivarin, act directly to form 2 and 1,1 hydrogen bonds with their target protein residues, mainly Cys172, Tyr435 with Cannabicyclol, Leu171 with Cannabigerolic acid, and tetrahydrocannabivarin. Cannabicyclol with Van der Waals interactions has been extensively documented, as have hydrogen bonding interactions with the primary reactive residues Gly58, Ser59, Tyr188, Ile198, Gln206, and Lys296 for Cannabicyclol, Gly57, Gly58, Ser59, Leu164, G569, G469, I169, Gly59, ISL264, ISL26, 1966, leu320, Met341, Tyr435 for Cannabigerolic acid, Gly40, THr43, Gly57, Gly58, Ser59, Tyr60, Phe343, Val294, Tyr398, Thr426, Gly434, Ala439 for tetrahydrocannabivarin, and Thr43, Gly57, Gly58, Ser59, Tyr188, Lys296, Thr426, Thr426, and Gly434 show Van der Waals interactions for Cannabidivarin. Several observed interactions are shown in Figure 1 and Table 7.

**Figure 1.**
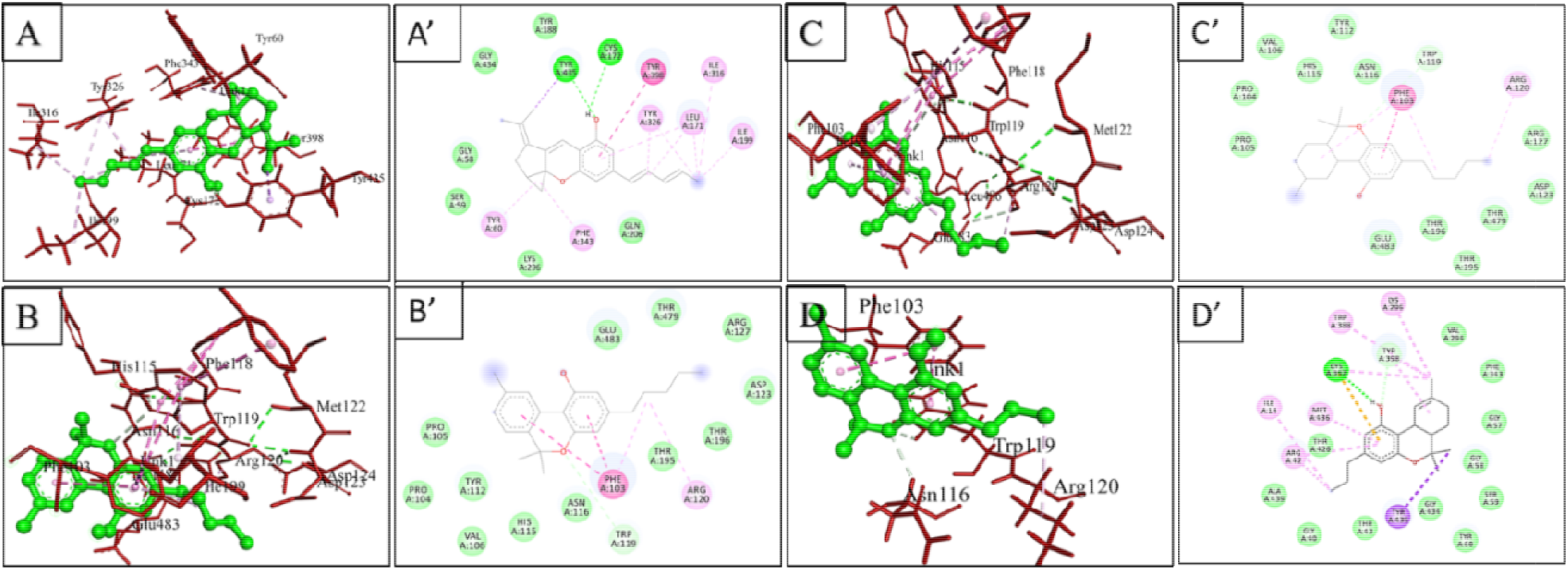
3D (left) and 2D (right) views of interacting residues of MAO-B with the four highest score compounds: (A, A’) (cannabicyclol), (B, B’) cannabinol, (C, C’) Tetrahydrocannabinol (D, D’) Cannabivarin

**Table 7.**
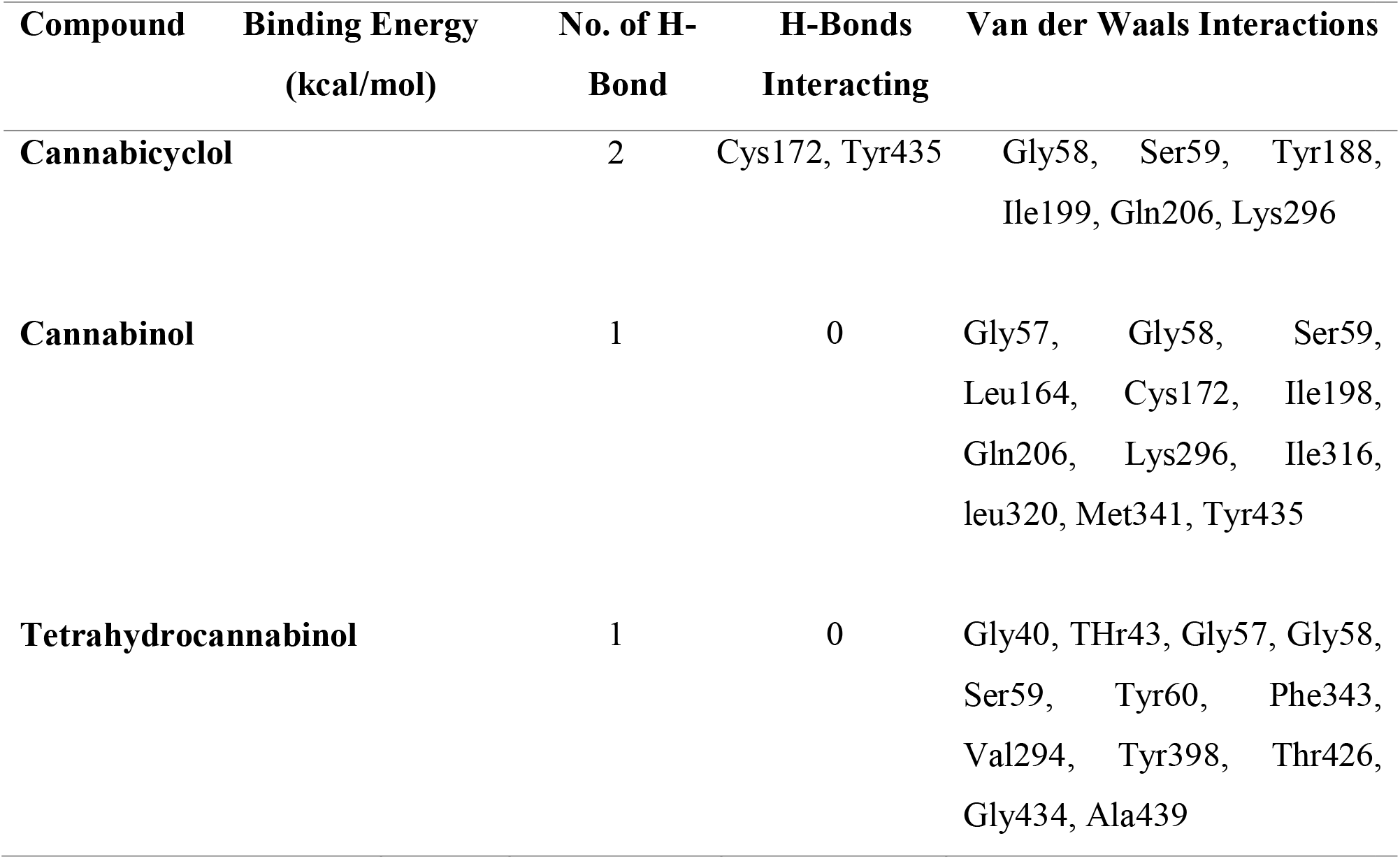
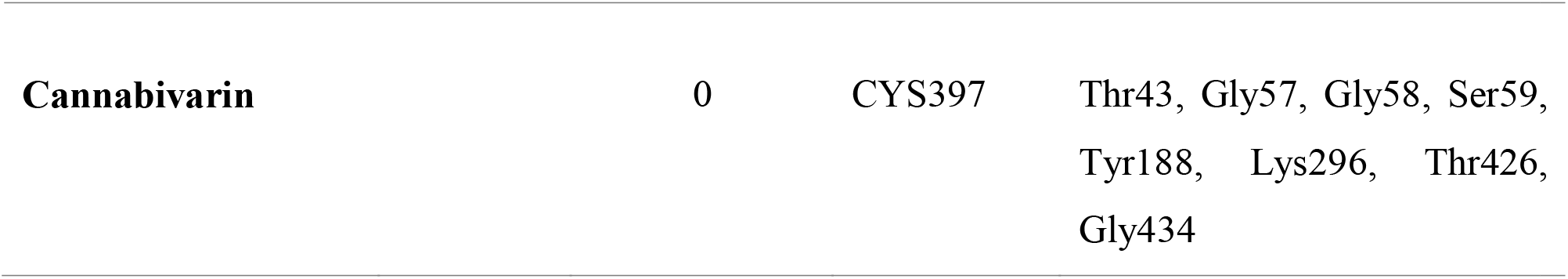
Show the interaction of various compounds with MAO-B protein residues.

### Molecular interaction of the Alpha-synuclein active residues with the highest docking score constituents

Cannabicyclol, Cannabitriol, Cannabicitran, and Tetrahydrocannabivarin have the highest docking scores of −6.4, −6.2, −6.1, and −6.0 against the target Alpha-synuclein. Residues of the protein alpha-synuclein (Val40, Lys43, and Lys45 for cannabitriol) represent three hydrogen bonds, and Val40 for Tetrahydrocannabivarin shows one hydrogen bond for each molecule. The compounds cannabicyclol and cannabicitran did not interact through hydrogen bonding. Van der Waals interactions are another well-observed type of interaction, in addition to hydrogen bonding. Residues Glu35, Gly36, Leu38, Tyr39, and Val49 show van der Waals interactions for cannabicyclol, Gly41, Thr44, and Val48 for cannabitriol; Glu35, Gly36, and Tyr39 for Cannabicitran; and residues Lys32, Glu35, Gly36, Gly41, and Lys45 interact with Tetrahydrocannabivarin by van der Waals interactions. In addition, some interactions were noticed and are shown in Figures 2 and Table 8.

**Figure 2.**
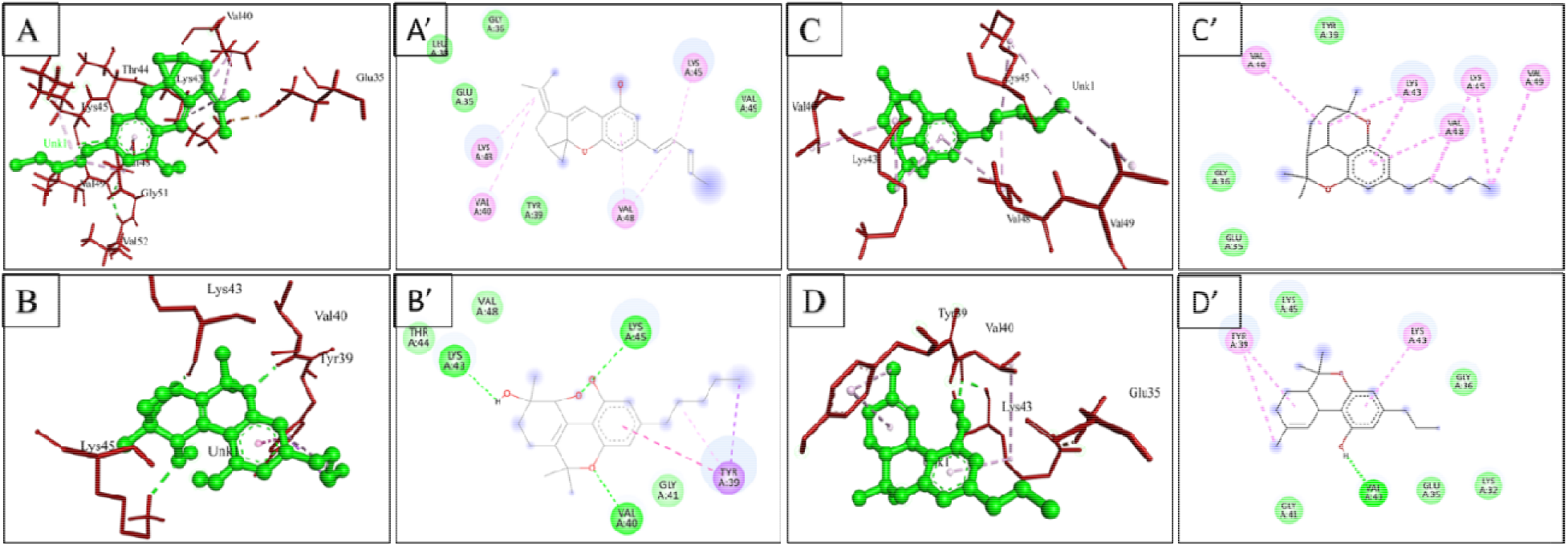
3D (left) and 2D (right) views of the interacting residues of Alpha-Synuclein with the four highest score compounds: (A, A’) (cannabicyclol), (B, B’) Cannabitriol, (C, C’) Cannabicitran, (D, D’) Tetrahydrocannabivarin

**Table 8.** Show the interaction of various compounds with Alpha-synuclein protein residues.

| Compound | Binding Energy<br>(kcal/mol) | No. of H-Bond | H-Bonds<br>Interacting | Van der Waals Interactions |
| --- | --- | --- | --- | --- |
| Cannabicyclol | -6.4 | 0 | 0 | Glu35, Gly36, Leu38,<br>Tyr39, Val49 |
| Cannabitriol | -6.2 | 3 | Val40, Lys43, Lys45 | Gly41, Thr44, Val48 |
| Cannabicitran | -6.1 | 0 | 0 | Glu35, Gly36, Tyr39 |
| Tetrahydrocannabivarin | -6.0 | 1 | Val40 | Lys32, Glu35, Gly36,<br>Gly41, Lys45 |

### Molecular interaction of the A2A active residues with the highest docking score constituents

Cannabicyclol, cannabinol, cannabitriol, and Cannabicitran had the highest docking scores at – −10.6, −9.7, −8.5, and −8.5 against the target A2A. The residue of protein A2A, Asn253, for Cannabicyclol, represents one hydrogen bond, and the same residue, Ans253, for cannabinol shows one hydrogen interaction with the A2A enzyme. The compounds cannabitriol and Cannabicitran did not interact through hydrogen bonding. Van der Waals interactions are another well-observed interaction type besides hydrogen bonding, residues Ala63, Thr88, Glu169, Asn181, Val186, Met270, and Tyr271 show van der Waals interactions for cannabicyclol, Ile66, Ser67, Ala81, Glu169, Tyr271, and His278 for cannabinol, Ile106, Arg107, Arg111, Arg206, Thr224, Leu225, Glu228, Arg1008, Ile1009, Gly1012, and Arg1014 for cannabitriol, and residues Ala59, Phe62, Ile80, Ala81, Val84, Thr88, Asn181, Asn253, Tyr271, and His278 interact with Cannabicitran by van der Waals interactions. In addition to this, some interactions have been noticed and may be shown in Figures 3 and Table 9.

**Figure 3.**
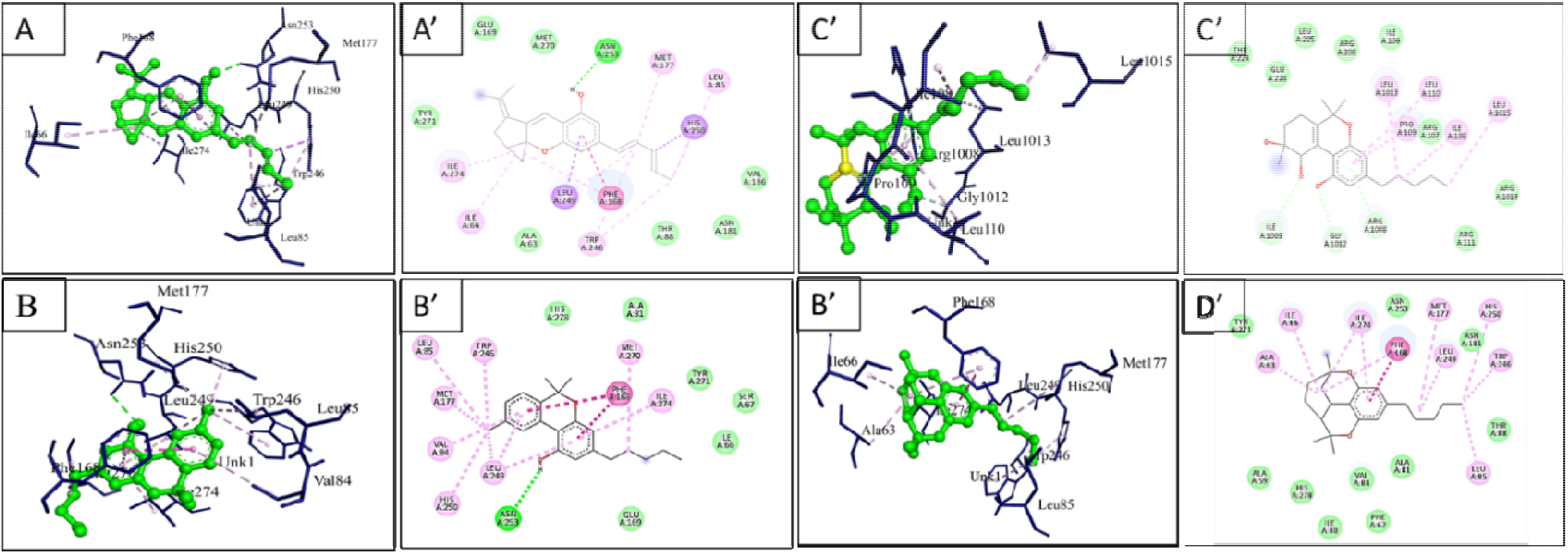
3D (left) and 2D (right) views of the interacting residues of A2A with the four highest score compounds: (A, A’) (cannabicyclol), (B, B’) cannabinol, (C, C’) cannabitriol, (D, D’) Cannabicitran

**Table 9.**
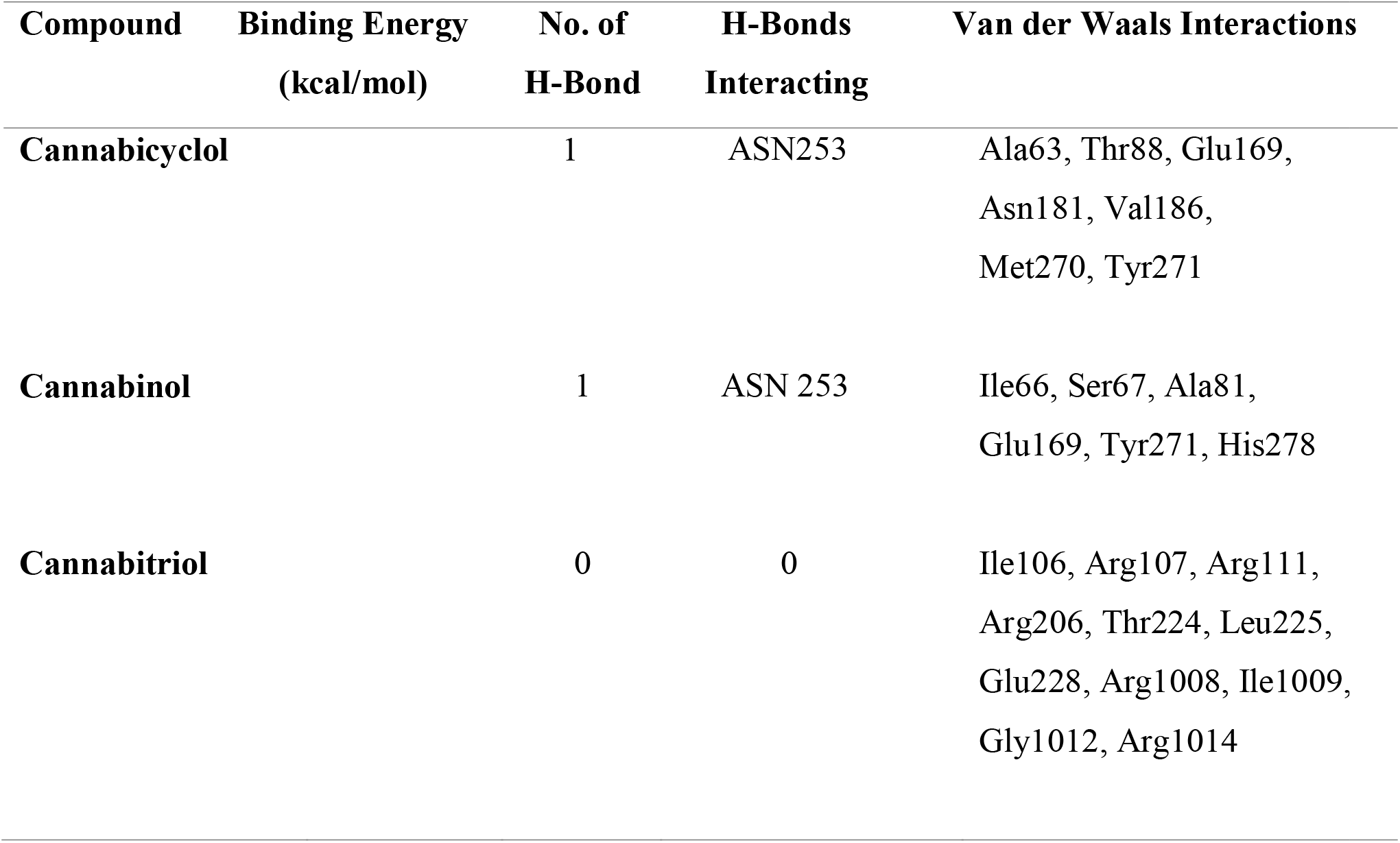
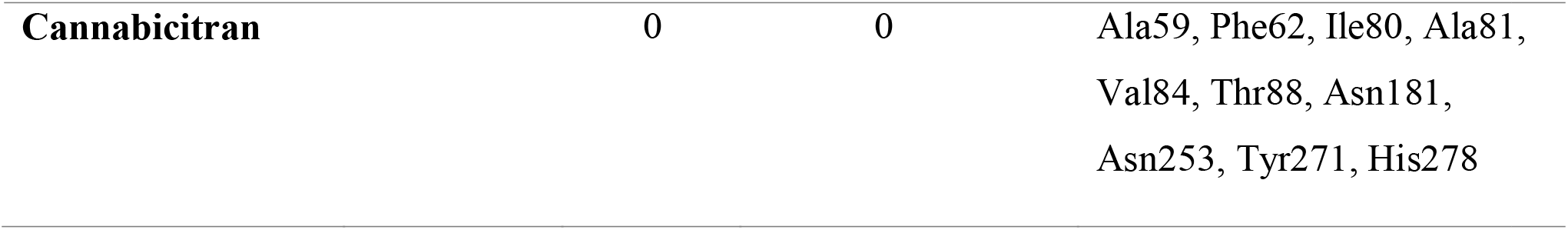
Show the interaction of various compounds with A2A protein residues.

### Molecular interaction of the COMT active residues with the highest docking score constituents

Ligands cannabicyclol, Cannabinol, cannabielsoin, and Cannabicitran had the highest scores of −6.4 kcal/mol, −6.4 kcal/mol, −6.1 kcal/mol, and −6.1 kcal/mol with the COMT protein, as shown below. Three substances, cannabicyclol, cannabielsoin, and Cannabicitran, act directly to form 1,1 hydrogen bonds with their target protein residues, mainly Asp195 with Cannabicyclol, Lys194 with cannabielsoin, and Cannabicitran. Van der Waals interactions have been extensively documented with the primary reactive residues Gly225, Ala226, Asp228 for Cannabicyclol, Gly57, Asp195, Pro224, Gly225, Ala226, Asp228 for Cannabinol, Gly40, THr43, Gly57, Gly58, Ser59, Tyr60, Phe343, Val294, Tyr398, Thr426, Gly434, Ala439 for Te Tyr118, Asp191, His192, Asp195 for cannabielsoin, and Trp88, His192, Trp193, Asp195, Asn220, Leu248, Glu249 show Van der Waals interactions for Cannabicitran. Moreover, other interactions were observed, as depicted in Figures 4 and Table 10.

**Figure 4.**
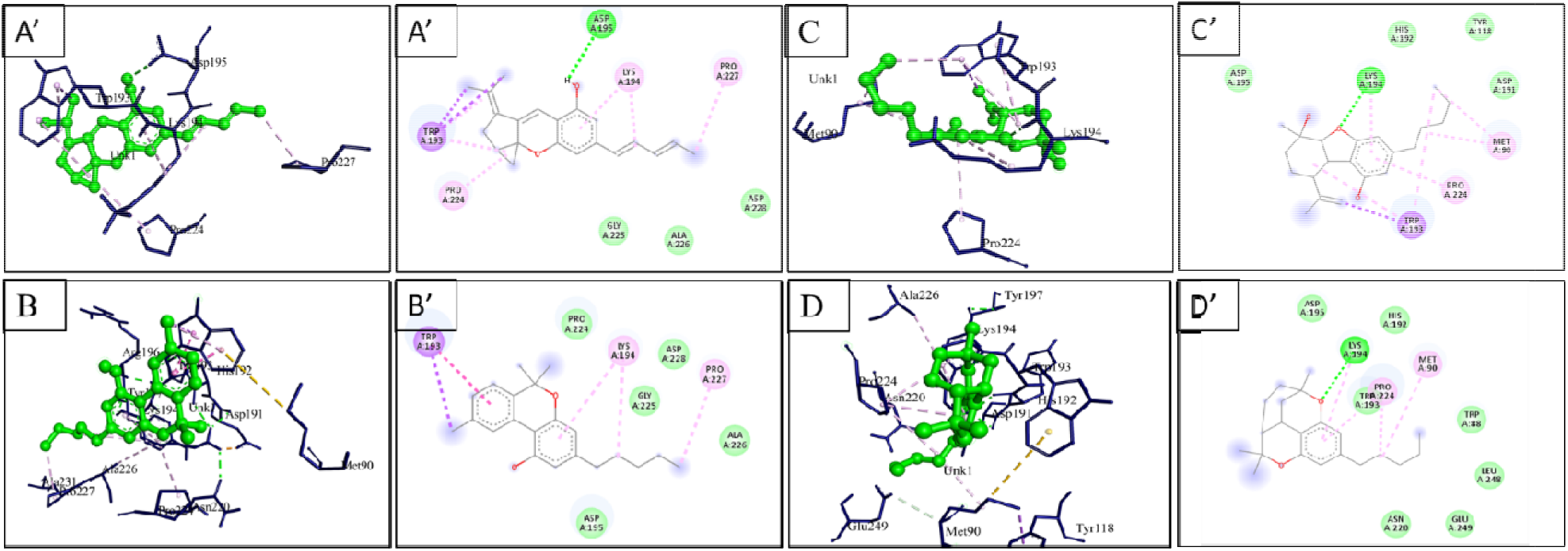
3D (left) and 2D (right) views of the interacting residues of COMT with the four highest score compounds: (A, A’) (cannabicyclol), (B, B’) cannabinol HD, (C, C’) cannabielsoin, (D, D’) Cannabicitran

**Table 10.**
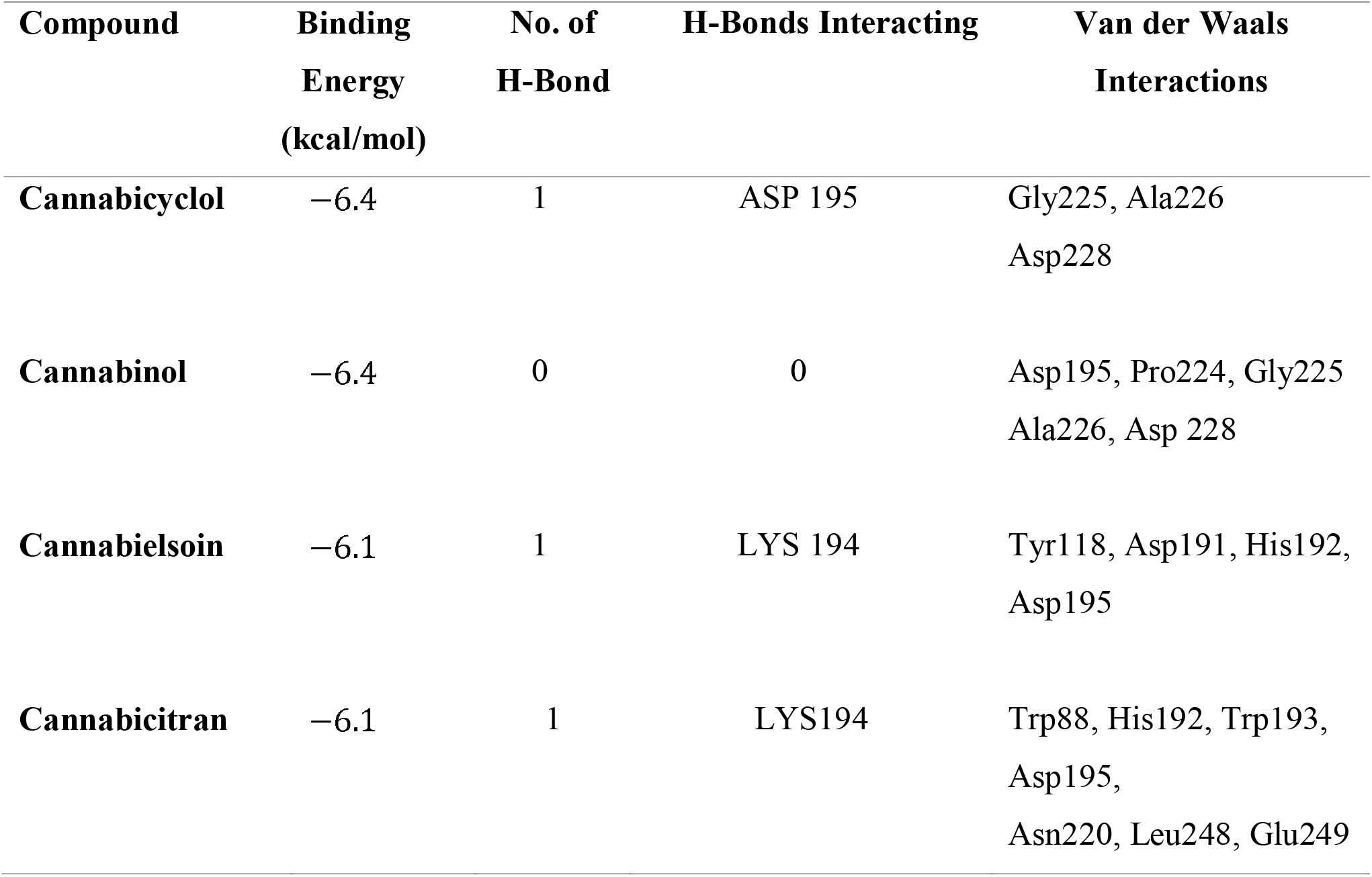
Show the interaction of various compounds with COMT protein residues.

### MD simulation

#### Root mean square deviation (RMSD)

For the first 23 ns of the simulation, the ligand had a relatively stable RMSD of 0.8 m. This demonstrates that the ligand retained its stable conformation during this time. However, the ligand underwent a sudden conformational change at 23 ns, which caused a higher RMSD value of 2.8 over the remaining 100 ns. This shows that after 23 ns, the ligand underwent another significant structural rearrangement or binding event with the protein, leading to a new stable conformation with higher structural divergence from the initial state.

According to the RMSD values, the protein may undergo structural changes during the first 23 ns, which fluctuate between 1.2 Å to 2.4 Å. This suggests that the protein is already flexible or that it is undergoing structural changes. However, the protein became more stable after 23 ns in the remaining 100 ns. This suggests that the protein settled into a stable shape after the initial fluctuations and maintained this stable conformation throughout the simulation. The RMSD graph provides crucial details on the behavior of the ligand and protein during MD simulation. After 23 ns, the ligand underwent a substantial conformational change with the protein, resulting in a new stable conformation. The protein’s structure initially fluctuates but eventually settles into a steady state that is maintained for the duration of the simulation (Figure 5 (**A**)).

**Figure 5.**
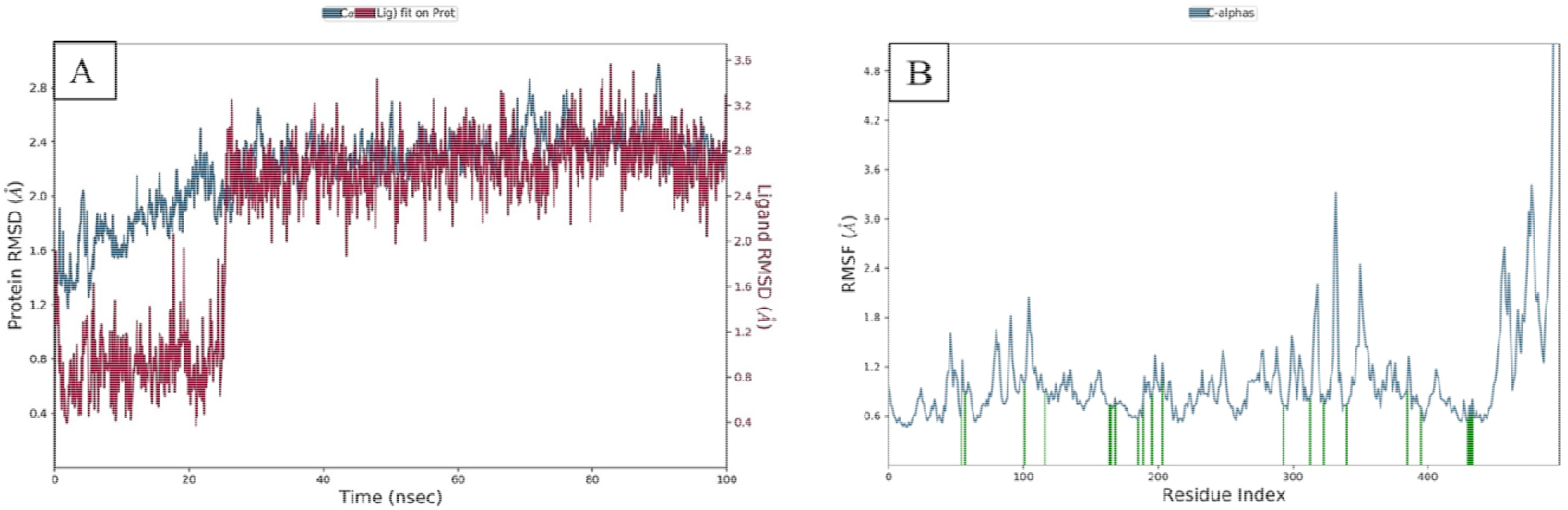
RMSD graph of Cα atoms of MAO-B protein with Cannabicyclol (A). RMSF graph of MAO-B residues (B)

### Root mean square fluctuation (RMSF)

According to the RMSF values in the 0.6 Å to 1.2 Å range, most protein residues typically undergo minor fluctuations during the simulation. Lower RMSF values (closer to 0.6) in a residue signify structurally stable protein areas that have not significantly changed from their initial conformation. These regions might correlate with stable secondary structures with high order, such as alpha helices or beta sheets. During the simulation, some residues or regions experienced more pronounced fluctuations, as indicated by the RMSF peaks higher than 1.2 Å. These more flexible portions of the protein may correspond to loops, turns, or other, more flexible locations. The higher RMSF values indicate that these regions are more dynamic and are probably subject to local movements or conformational changes throughout the simulation (Figure 5 **B**).

### Ligand-Protein contact

The protein-ligand contact analyses of the studied complexes were estimated throughout the 100 ns of MD simulations and analyzed using the contact histogram model. A total of 25 protein residues were in contact with cannabicyclol, from Gly 57 to Met436. The generated histograms show H-bond residues consisting of Cys172, Tyr188, Tyr398, Gly434, and Tyr435 with Cannabicyclol. Our results demonstrated the presence of a variety of interactions between the active site residues of MAO-B and the tested ligand, cannabicyclol, including H-connections, Water bridges, and hydrophobic and ionic bonds. H-bonds are divided into sub-types essential for ligand binding and drug selectivity, such as backbone acceptor and backbone donor, side-chain donor, and side-chain acceptor. However, some residues formed hydrophobic and non-specific interactions involving π-cation, π–π, and hydrophobic amino acid ligands (Figure 6 (**A**)). The top panel represents the total number of interactions between the protein forms and ligand. The amino acids that interacted with the ligand in each trajectory frame are shown in the bottom panel. The scale to the right of the figure indicates that specific residues made more than one contact with the ligand, as indicated by the darker orange shade in Figure 6 (**B**).

**Figure 6.**
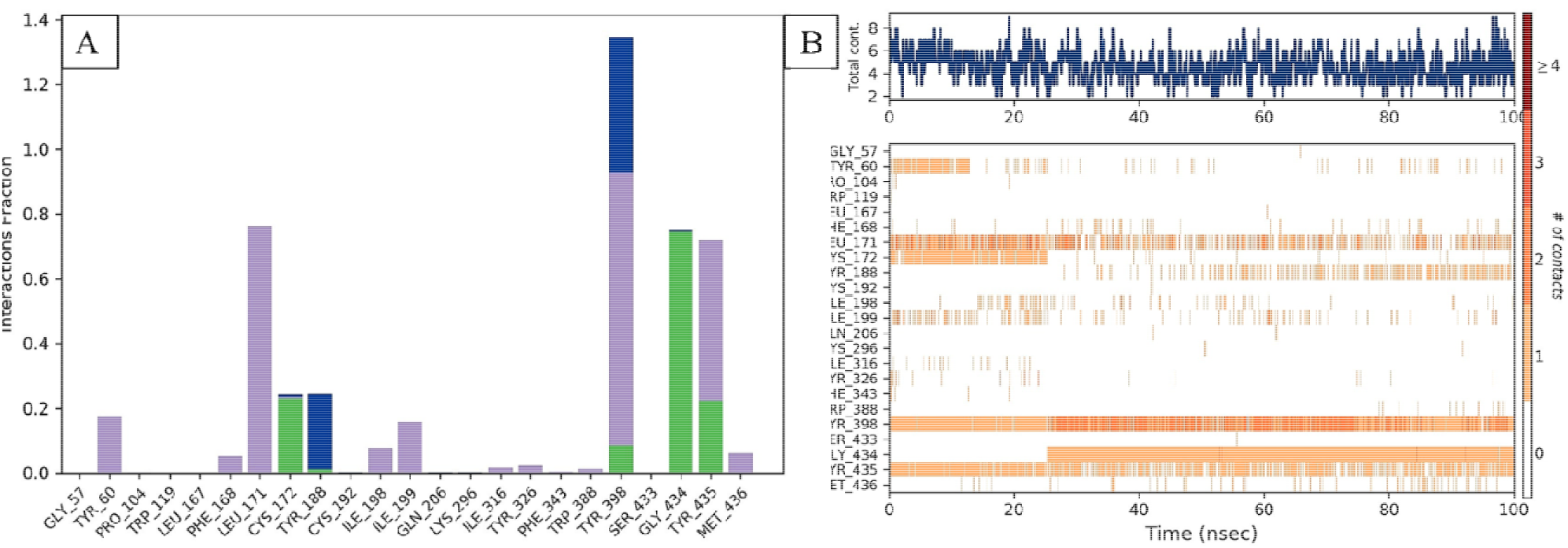
Total number of contacts of MAO-B protein bound to cannabicyclol (A). A timeline illustrating the interaction and contacts (H-bonds, hydrophobic, ionic, and water bridges) (B)

### Ligands properties analysis

Its characteristics were studied in depth to demonstrate the geometrical and structural changes of the ligand in a 100 ns simulation. RMSD, radius gyration (rGyr), molecular surface area (MolSA), accessible surface area (SASA), and polar surface area (PSA) of the ligand were determined as functions of time. The RMSD study represents slight variation in interaction with MAO-B for ligand cannabicyclol (approximately 0.3 to 0.9 Å, equilibrium ∼ 0.3 Å). The measured rGyr values varied from 4.08 to 4.32 Å (equilibrium ∼ 4.15 Å). The MolSA values ranged from 309 to 318 Å2 (equilibrium, 280 2). Ultimately, the observed SASA values fluctuated from 0 to 60 Å2 during the simulation. A slight Variation in PSA values was observed for this complex, and equilibrium was established at 57 Å2. In this scenario, the ligand demonstrated little fluctuation and maintained equilibrium, indicating remarkable stability in the active pocket of MAO-B (Figure 7).

**Figure 7.**
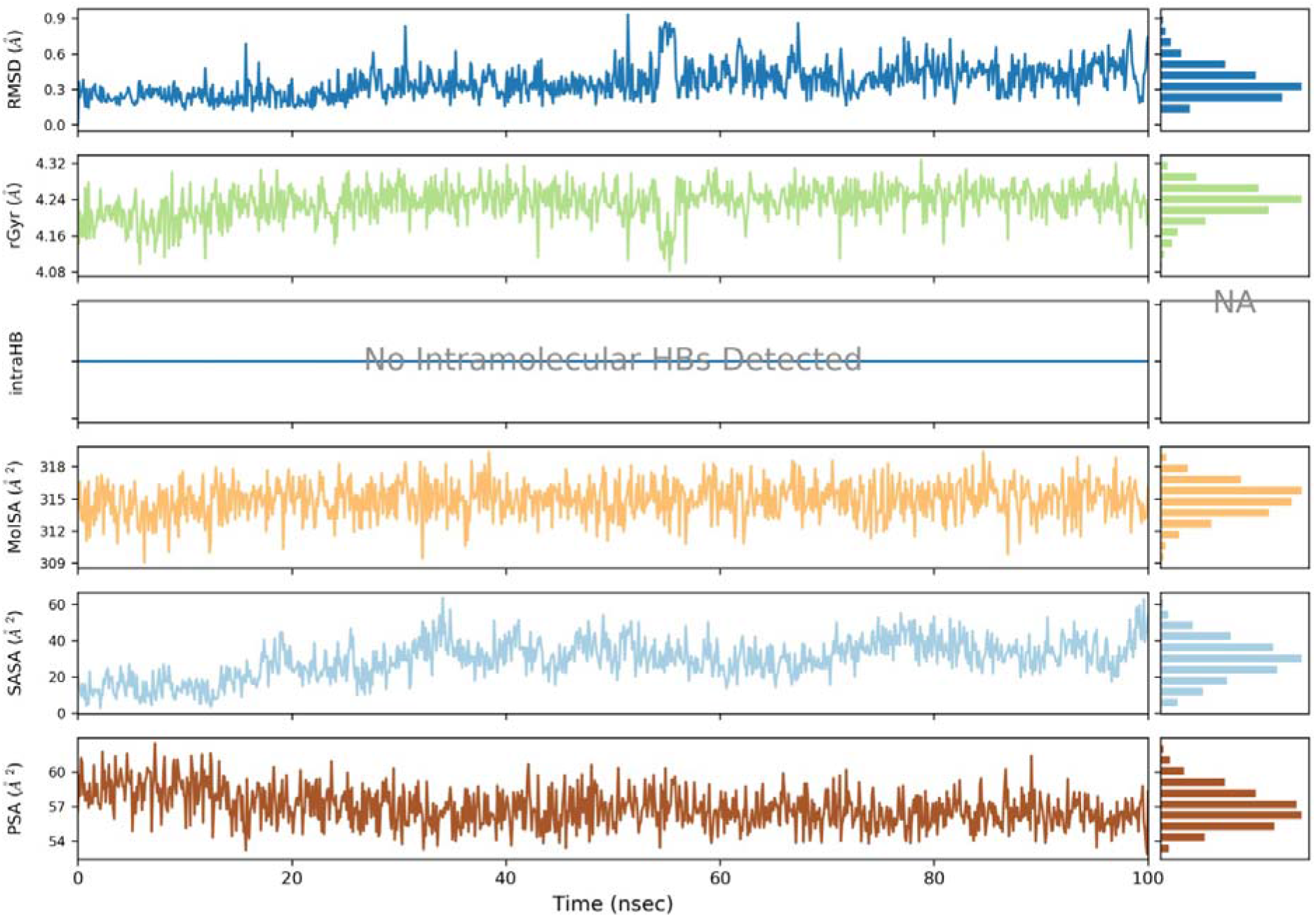
Properties of Cannabicyclol in complex with MAO-B during 100 ns Simulation

## Discussion

The present study aimed to perform molecular docking experiments to investigate the therapeutic potential of certain cannabis constituents for the treatment of Parkinson’s. In this study, we identified four potential inhibitors of all Parkinson’s disease-associated proteins. Our results indicate that cannabicyclol exhibited the best docking scores for all target proteins Alpha-synuclein, MAO-B, A2A, and COMT, illustrating its high affinity and potential therapeutic efficacy in these neurodegenerative conditions. Many health benefits of cannabis constituents have been observed, including their ability to protect the brain from conditions, such as Parkinson’s disease. High docking scores for cannabicyclol indicate that this drug inhibitor may reduce the symptoms of Parkinson’s by targeting dopamine pathways. As a precursor molecule, cannabicyclol may be a suitable inhibitor of dopamine reuptake and a MAO-B inhibitor to increase dopamine levels [75]. These results are consistent with earlier research showing that cannabinoids may reduce neuroinflammation, oxidative stress, and neuronal damage related to Parkinson’s [76].

Cannabicyclol is a non-psychoactive and non-toxic cannabinoid. This is a result of the degradation of cannabichromene, a phytocannabinoids. According to the current research, cannabicyclol can be considered non-psychoactive because it has no recognized physiological effects. Cannabicyclol may have antibacterial, anti-inflammatory, neurogenic (boost brain cell proliferation), and antiemetic qualities because of its structural resemblance to Cannabichromene and Cannabinol [77–79].

Cannabinoids have also been linked to several beneficial pharmacological benefits but may also have unpleasant consequences. On the one hand, cannabinoids have been linked to many advantageous pharmacological effects, but on the other, they have been linked to toxic and unfavorable side effects. Recent research indicates that the number and age of consumers also directly affect health [80]. Hence, research on pathways associated with PD and related enzyme regulatory studies is essential. It has been demonstrated that cannabinoids reduce oxidative stress and excitotoxicity and control dopamine levels [81].

In this study, cannabicyclol was found to be a significantly more appropriate compound [82]. These results indicate that cannabicyclol can bind to the alpha-synuclein active site and inhibit protein self-association to provide Parkinson’s disease treatment. As alpha-synuclein is a hallmark of Parkinson’s disease, it plays a crucial role in diagnosing and causing PD disorder [83]. The findings of the present study support in silico molecular docking investigations of certain cannabis compounds and specific pharmacological ingredients against the MAO-B enzyme [84]. These findings demonstrate that, in comparison to the established medications Deprenyl, Rasagiline, and selegiline, the leads, in particular Cannabicyclol, Cannabitriol, Cannabivarin, Cannabinol, and Cannabicitran, showed higher binding affinities and interactions with the MAO-B enzyme [85]. Cannabis compounds can be used as medicinal agents to create therapeutically useful MAO inhibitor [86].

Surrounding the COMT’s work area are two large hydrophobic areas, just next to the methylation site, making the space around it smaller and tighter [87]. The amine and carboxylic groups linked to the sp3-carbon of L-DOPA hardly fit between the targeted residues, which tends to cause the catechol substituent to move in the opposite direction, where the ionized groups are heavily solvated [88]. These steric restrictions help to fill out catechol-binding modes that favor O-methylation at the meta (MO) position [89]. Therefore, despite being strictly qualitative, the current results provide a different explanation for the observed COMT regioselectivity with L-DOPA [90]. These findings demonstrate that, in comparison to the established medications entacapone and tolcapone, the leads, in particular Cannabicyclol, Cannabitriol, Cannabinodiol, Cannabinol, and Cannabicitran, showed greater binding affinity and interactions with COMT [91]. Cannabis compounds can be used as medicinal compounds to create therapeutically useful COMT inhibitor [11].

The A2A receptor-drug complex may have a higher binding affinity owing to hydrogen bonds. The A2A receptor’s active amino acid residues show good attraction for cannabis compounds, including cannabicyclol, cannabitriol, cannabivarin, cannabinol, and Cannabicitran [92]. The significant connection between adenosine and dopamine receptors in the brain has made the adenosine A2A receptor a key non-dopaminergic treatment target for Parkinson’s [93]. According to research, A2A receptor antagonists have shown the potential to reduce the use of Parkinson’s medications. Other drugs targeting non-dopaminergic systems include ASN inhibitors, MAO-B inhibitors, dopamine agonists, COMT inhibitors, and A2A inhibitors [94].

The physicochemical properties of the cannabis constituents, including their polar surface area, moderate molecular weight, and lipophilicity, support their advantageous drug-like effects. These characteristics increase the possibility of effective drug delivery, bioavailability, and engagement of target proteins. Furthermore, cannabis constituents can enter the brain and exhibit therapeutic effects, as shown by the expected ADME/T qualities, which include strong GI absorption, BBB permeability, and CNS permeability [95].

Through molecular docking and MD simulations, we examined the therapeutic potential of the cannabis ingredient cannabicyclol in the treatment of Parkinson’s disease. The target proteins Alpha-synuclein, MAO-B, A2A, and COMT showed cannabicyclol with the best docking scores, indicating high affinity. Based on the low binding energy and hydrogen bonding, the MD simulations showed that cannabicyclol and MAO-B had a stable and durable association. These results highlight the potential inhibition of MAO-B activity by cannabicyclol. Validating these results and investigating the therapeutic uses of cannabis in neurodegenerative disorders requires further investigation.

This in silico research shows that some newly discovered cannabis compounds have more inhibitory effects on these enzymes, which may be better than existing ones. Our research offers molecular docking evidence in favor of cannabicyclol for the treatment of Parkinson’s disease. According to these researchers, cannabicyclol should be further studied as a potential therapeutic option for various neurological disorders because of its favorable physicochemical features, expected ADME/T properties, and interactions with target proteins. To further our understanding and establish the clinical efficacy of cannabis in managing Parkinson’s disease, future research, including in vitro and in vivo investigations, as well as clinical trials, is required.

## Conclusion

In silico approaches indicated that cannabis constituents have potential neuroprotective properties. Compared to conventional medicines against PD, cannabis constituents show greater potency in inhibiting the enzymes A2AR, COMT, MAO-B, and alpha-synuclein, which contribute to Parkinson’s disease. This in silico research suggests that cannabicyclol has greater efficiency in inhibiting all these enzymes, which act as culprits in Parkinson’s disease. Furthermore, in vitro and in vivo research is required to increase the effectiveness of these cannabis constituents and to better understand the pathways that inhibit the enzymes associated with Parkinson’s conditions.

